# The *Candida albicans* ζ-crystallin homolog Zta1 promotes resistance to oxidative stress

**DOI:** 10.1101/2023.09.05.556406

**Authors:** Rafael M. Gandra, Chad J. Johnson, Jeniel E. Nett, James B. Konopka

**Affiliations:** Department of Microbiology and Immunology, Stony Brook University, Stony Brook, New York, United States of America; University of Wisconsin-Madison, Department of Medicine; University of Wisconsin-Madison, Department of Medical Microbiology & Immunology

**Keywords:** *Candida albicans*, ζ-crystallin, Zta1, oxidative stress, quinone, quinone reductase, flavodoxin like protein

## Abstract

The fungal pathogen *Candida albicans* is capable of causing lethal infections in humans. Its pathogenic potential is due in part to the ability to resist various stress conditions in the host, including oxidative stress. Recent studies showed that a family of four flavodoxin-like proteins (Pst1, Pst2, Pst3, Ycp4) that function as quinone reductases promotes resistance to oxidation and is needed for virulence. Therefore, in this study Zta1 was examined because it belongs to a structurally distinct family of quinone reductases that are highly conserved in eukaryotes and have been called the ζ-crystallins. The levels of Zta1 in *C. albicans* rapidly increased after exposure to oxidants, consistent with a role in resisting oxidative stress. Accumulation of reactive oxygen species was significantly higher in cells lacking *ZTA1* upon exposure to quinones and other oxidants. Furthermore, deletion of *ZTA1* in a mutant lacking the four flavodoxin-like proteins, resulted in further increased susceptibility to quinones, indicating that these distinct quinone reductases work in combination. These results demonstrate that Zta1 contributes to *C. albicans* survival after exposure to oxidative conditions, which increases the understanding of how *C. albicans* resists stressful conditions in the host.

**IMPORTANCE:** *Candida albicans* is an important human pathogen that can cause lethal systemic infections. The ability of *C. albicans* to colonize and establish infections is closely tied to its highly adaptable nature and capacity to resist various types of stress, including oxidative stress. Previous studies showed that four *C. albicans* proteins belonging to the flavodoxin-like protein family of quinone reductases are needed for resistance to quinones and for virulence. Therefore, in this study we examined the role of a distinct type of quinone reductase, Zta1, and found that it acts in conjunction with the flavodoxin-like proteins to protect against oxidative stress.

## INTRODUCTION

*Candida albicans* is a commensal organism that can grow on the skin and mucosa of healthy individuals, but under certain circumstances it can overgrow and spread to cause severe mucosal infections or lethal systemic infections (1). In order to be an effective pathogen, *C. albicans* must resist stressful conditions to survive in the host, such as elevated temperature, oxidative and nitrosative stress, antimicrobial peptides, and regulation of micronutrients, such as iron depletion or copper toxicity (2-5). One important type of stress encountered in vivo is caused by reactive oxygen species (ROS), which damage DNA, lipids and proteins and lead to cell death (6). *C. albicans* uses an array of antioxidant mechanisms to deal with ROS, such as superoxide dismutase, catalase, thioredoxin and glutathione (1).

Recently, a new antioxidant pathway was described in *C. albicans* involving four flavodoxin-like proteins (FLPs; Pst1, Pst2, Pst3 and Ycp4) that are important for resisting oxidative stress, preventing lipid peroxidation, and for virulence in mice (7, 8). The FLPs belong to a large family of NADPH:quinone oxidoreductases that are present in many organisms, including bacteria, fungi, and plants, but are not conserved in mammals (9). FLPs are enriched in plasma membrane eisosome domains where they are thought to protect the *C. albicans* plasma membrane by reducing ubiquinone to ubiquinol so that it can reduce reactive oxygen species (7). FLPs are also important for resisting oxidative stress caused by treatment of *C. albicans* with a variety of smaller quinone molecules (7, 8). FLPs are thought to therefore play a critical role in protecting cells from quinones that are created as secondary metabolites, or in many cases are used by plants and insects as defense molecules (10-12). Due to their redox-active nature, quinones can cause oxidative stress by undergoing cycles of oxidation and reduction, which can consume reducing agents such as glutathione and NADPH. In addition, FLPs carry out a two-electron reduction of quinones, which prevents the formation of dangerous semiquinone radicals that interact with oxygen to form superoxide anion radicals, leading to cell damage (10, 11).

In addition to FLPs, many organisms contain members of a distinct group of quinone oxidoreductases (QORs) that belong to the medium-chain dehydrogenase/reductase (MDR) superfamily (13, 14). The first QOR of this group to be described was the mammalian ζ- crystallin, a major protein of the eye lens that was observed to also be present in other cell types where it is capable of reducing quinones using NADPH as a cofactor (15-18). However, unlike the FLPs, it is believed that these enzymes catalyze a one-electron reduction of quinones (16). A ζ-crystallin homologous protein called Zta1 has been described in *Saccharomyces cerevisiae* and *Pichia pastoris*, and it has been suggested that it acts as a NADPH:quinone oxidoreductase that protects cells from oxidative stress (17, 18). However, little is known about it since data are limited for *S. cerevisiae* Zta1 and *C. albicans* Zta1 has not been studied previously. Therefore, in this study we examined the production of Zta1 and its function in protecting *C. albicans* from oxidative stress. In addition, a mutant lacking *ZTA1* as well as the four flavodoxin-like proteins (*zta1Δpst1 Δpst2 Δpst3 Δycp4Δ)* was constructed to determine determine a cell lacking all 5 QOR genes. These data indicate that Zta1 acts in combination with FLPs to protect *C. albicans* from oxidative stress.

## RESULTS

### Zta1 proteins are highly conserved between *C. albicans* and other fungi

The medium-chain dehydrogenase/reductase family includes the QOR known as ζ- crystallin that was first discovered in the lenses of camels and guinea pigs, but was subsequently found in a wide range of eukaryotic cells (19). Interestingly, a protein with homology to ζ-crystallin was identified in fungi. The crystal structure of the *Saccharomyces cerevisiae* ζ-crystallin, termed Zta1, has been described and shows that it is structurally similar to mammalian ζ-crystallin proteins (20). Comparison of the Zta1 proteins from *S. cerevisiae* and *C. albicans* revealed a high degree of sequence identity (56.8%) and there was 100% conservation of residues implicated in catalytic activity (Fig. 1). Although the human homolog Cryz has low overall sequence identity to Zta1 (∼26%), the active site residues are mostly conserved. Therefore, *C. albicans* Zta1 belongs to the highly conserved family of ζ-crystallin proteins.

**Fig 1.**
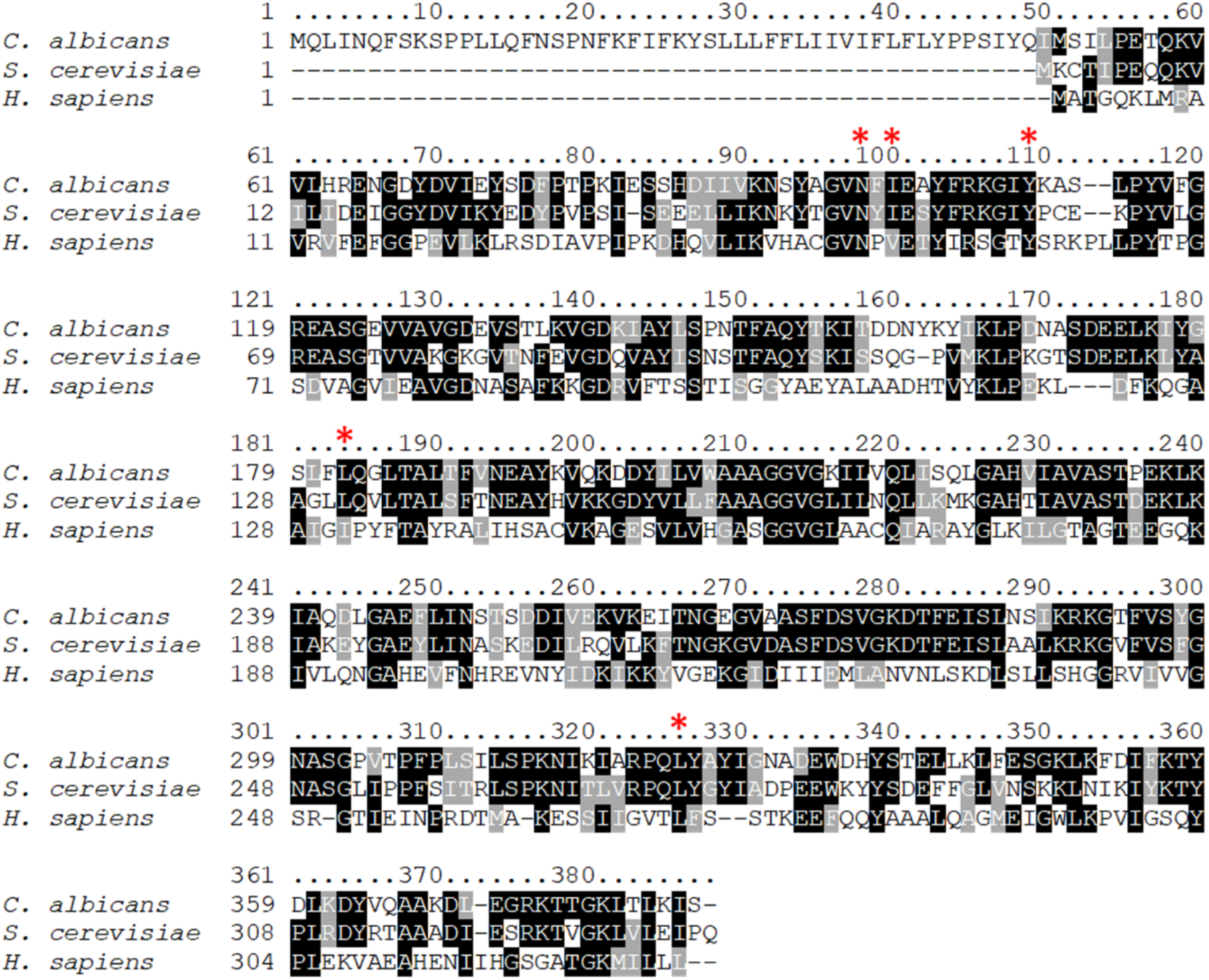
Conservation of Zta1 protein sequence. ClustalW alignment of the amino acid sequences of Zta1 from *Candida albicans, Saccharomyces cerevisiae* and the human homologue *Cryz*. Identical residues are highlighted in black and similar residues are shaded in gray. The red asterisk indicates Zta1 active site residues. A high degree of amino acid similarity was found at these sites.

### Zta1 is localized to the cytoplasm and induced by oxidation

To examine the subcellular localization and production of Zta1, a triple GFP tag (3xGFP) was fused to the 3’ end of the *ZTA1* open reading frame. Analysis of log-phase cells by fluorescence microscopy revealed that Zta1-3xGFP localized to the cytoplasm. The basal level of Zta1-3xGFP was low in cells grown in standard medium, but the signal was induced by the treatment with several different quinones including p-benzoquinone (BZQ) (4-fold), 2-tert-butyl-1,4-benzoquinone (TBBQ) (1.6-fold), and menadione (MEN) (2.5-fold). (Fig. 2A, D). Interestingly, H_2_O_2_ also significantly increased Zta1 production (3.5-fold), indicating that *ZTA1* expression is induced by oxidative stress, not just by quinones. Concentrations as low as 10 µM BZQ were sufficient to achieve a 3-fold increase in Zta1-3xGFP production (Fig. 2B, E). Treatment with 100 µM BZQ resulted in a reduction of Zta1-3xGFP induction, probably due to toxicity. *ZTA1* was induced quickly after exposure to BZQ. The relative level of Zta1-3xGFP increased by 2-fold and 3.1-fold increase after 30 and 60 minutes, respectively (Fig 2C, F).

**Fig. 2.**
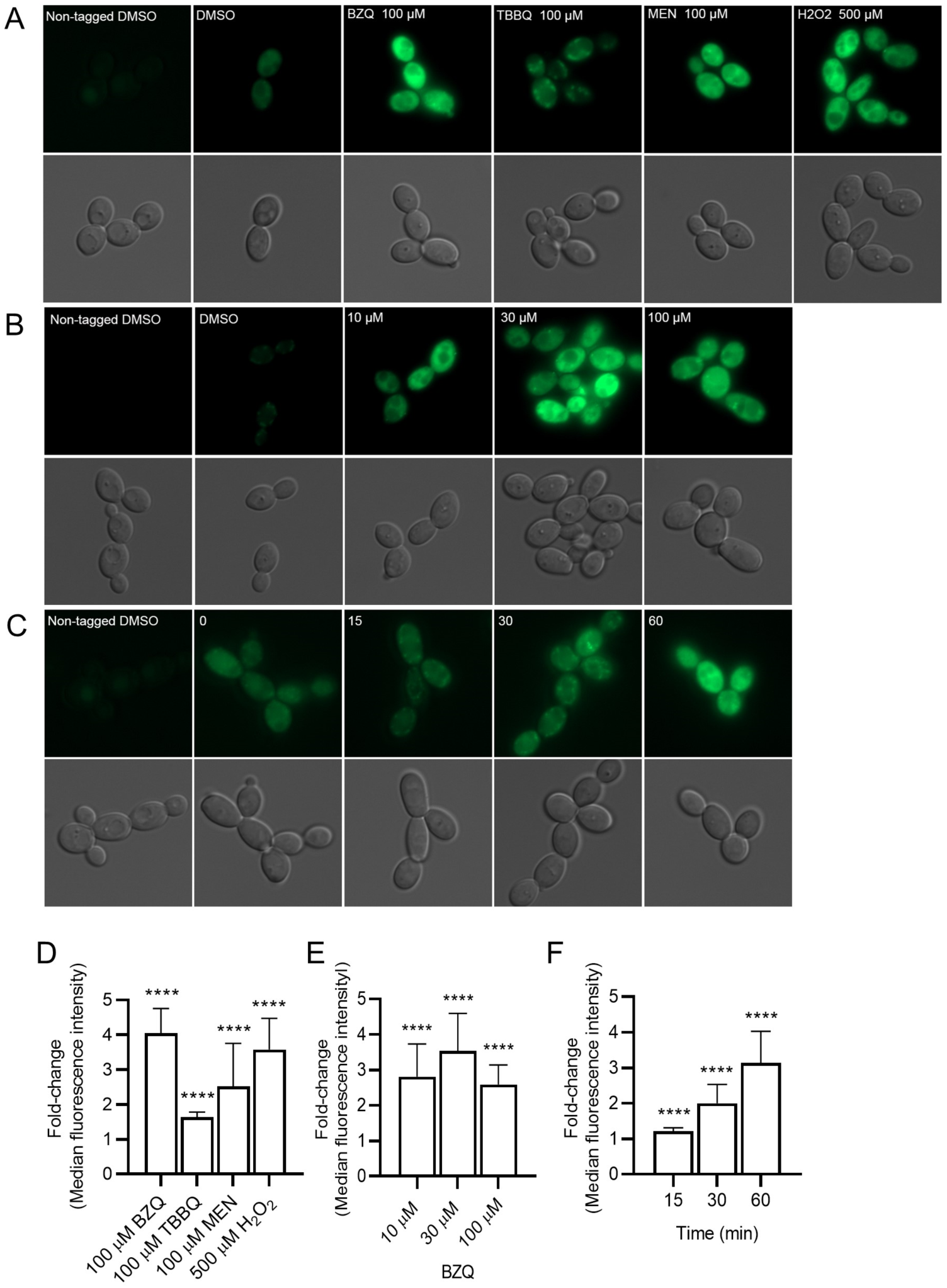
Fluorescence microscopy of *C. albicans* cells producing Zta1. (A) *C. albicans ZTA1-3xGFP* cells were incubated with 100 µM of p-benzoquinone (BZQ), 2-tert-butyl-1,4-benzoquinone (TBBQ), Menadione (MEN) and 500 µM of hydrogen peroxide (H_2_O_2_) at 30°C for 1 h, washed and then imaged by fluorescence microscopy. (B) *C. albicans ZTA1-3xGFP* cells were treated with different concentrations of p-benzoquinone (10, 30 and 100 µM) for 1 h and then analyzed. (C) cells were treated with 100 µM of p-benzoquinone and imaged after the indicate minutes of incubation. (D-F) Median fluorescence intensity of *ZTA1-3xGFP* cells treated with (D) the indicated compounds for 1 h, (E) different concentrations of p-benzoquinone for 1 h and (F) 100 µM p-benzoquinone for the indicated times. The bars represent the average fold-change of 3 independent assays performed on different days. *t* tests were performed comparing each condition to the control cells treated with DMSO. ****, *P* ≤ 0.0001.

Since Zta1*-*3xGFP was localized to the cytoplasm, Western blot analysis was carried out to determine whether the expected fusion protein was produced or if free GFP was being proteolytically clipped off. A band of ∼118 kDa was detected with anti-GFP antibodies, indicating the full length Zta1-3xGFP fusion protein was produced. Only a weak signal was detected at the expected position for free GFP (∼26.7kDa), confirming that Zta1-3xGFP is cytoplasmic.

Western blot analysis also detected increased production of Zta1-3xGFP after 1 hour induction with BZQ (2.3-fold), MEN (2.8-fold) and H_2_O_2_ (2.6-fold) (Fig. 3A). However, treatment with 100 µM TBBQ led to lower Zta1 induction (0.6-fold), probably due to greater toxicity. This was corroborated by results showing that TBBQ induced significantly more cell death after 1 h incubation than the other compounds (Fig. 3D). Additional analyses confirmed that Zta1 is induced by low concentrations of BZQ (2.5-fold at 10 µM; 3.9-fold at 30 µM) but starts to decrease at 100 µM (1.8-fold) (Fig. 3B). A rapid increase in Zta1 levels was observed after 15 minutes (3.6-fold), increasing to 6-fold after 30 minutes and by more than 10-fold after 1 hour of BZQ treatment (Fig. 3C). These data demonstrate Zta1 is rapidly induced by oxidizing agents.

**Fig. 3.**
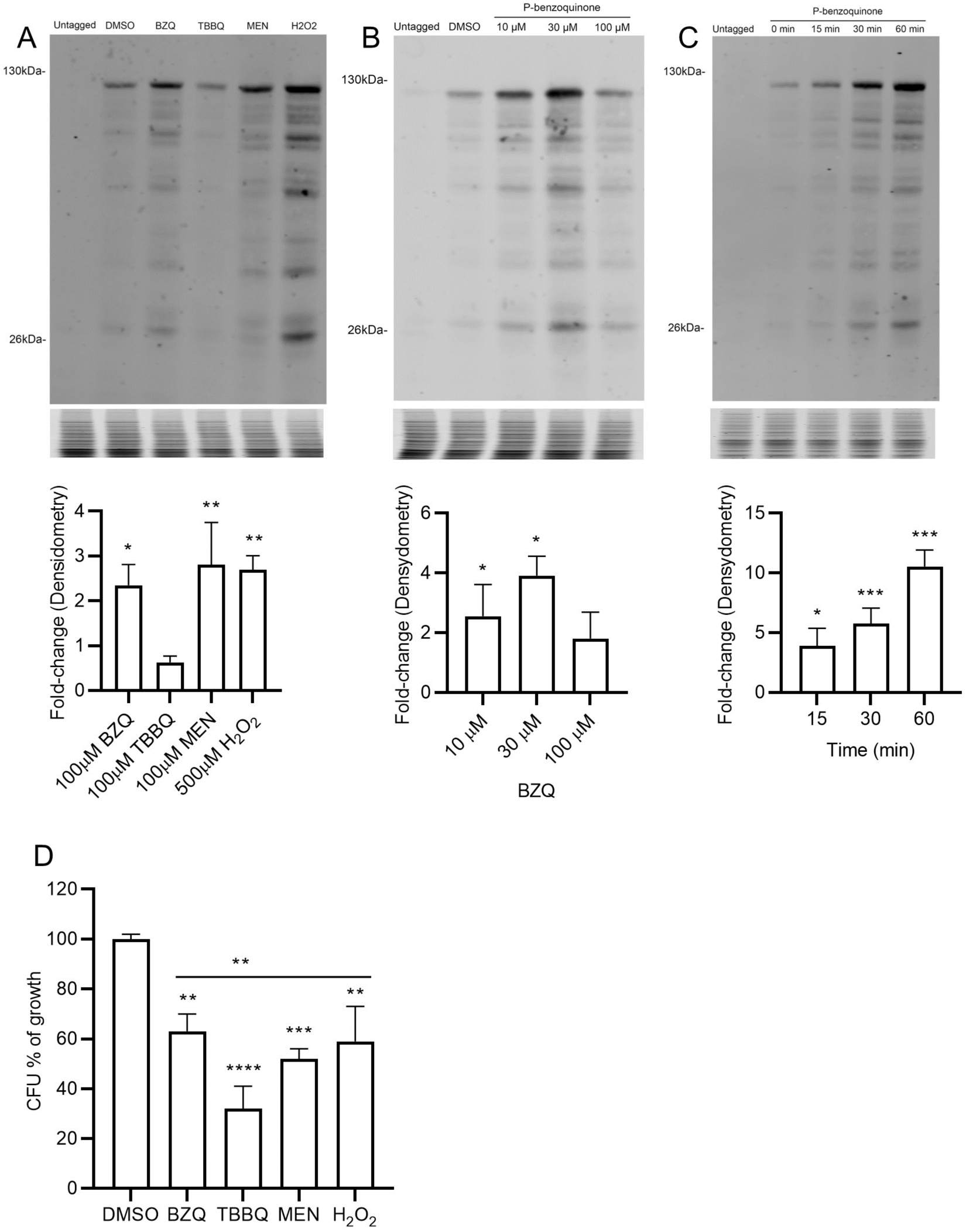
Zta1 is induced by quinones and oxidation. Western blots comparing Zta1-3xGFP production in *C. albicans* exposed to (A) the indicated oxidants for 1 h, (B) different concentrations of p-benzoquinone for 1 h, or (C) 100 µM of p-benzoquinone for the indicated times. Cells exposed to DMSO were used as a control. Western blots probed with anti GFP antibodies are shown on the top, Coomassie Blue stained gels of the protein samples shown below were used as a loading control. The bar graphs underneath showing quantitation of the results was performed using Image Studio software, normalized to Coomassie-stained gels. The results represent the average of three independent experiments performed on different days. (D) CFU assay of Zta1-3xGFP tagged cells treated with 100 µM of BZQ, TBBQ, MEN and 500 µM of H_2_O_2_ for 1 h. Results represent averages from three independent experiments performed on different days. Multiple *t* tests were performed comparing each condition to the DMSO-treated control. **, *P* ≤ 0.01; ***, *P* ≤ 0.001, **** *P* ≤ 0.0001. *t* tests were also performed comparing TBBQ with the other conditions, **, *P* ≤ 0.01.

### Zta1 prevents accumulation of reactive oxygen species

The ability of Zta1 to protect against ROS was examined using the fluorescent probe H_2_DCFDA, which has been previously used to quantify ROS accumulation in *Candida* cells (21). Wild type and *zta1Δ/Δ* mutant cells treated with 30 µM of the quinones BZQ and TBBQ did not lead to detectable accumulation of ROS using H_2_DCFDA (Fig. 4A, D). However, both quinones significantly induced ROS accumulation at 100 µM in the *zta1Δ/Δ* mutant, which was abolished in the complemented strain *zta1+* (Fig. 4B, C, E, F). The median fluorescence intensity of *zta1Δ/Δ* cells that were incubated with TBBQ was significantly higher when compared with cells exposed to BZQ (*t* test *P=* 0.021). These data indicate Zta1 can reduce ROS accumulation caused by quinones and protect *C. albicans* from oxidative damage, and that TBBQ induces higher production ROS than does BZQ in the *zta1Δ/Δ* cells.

**Fig. 4.**
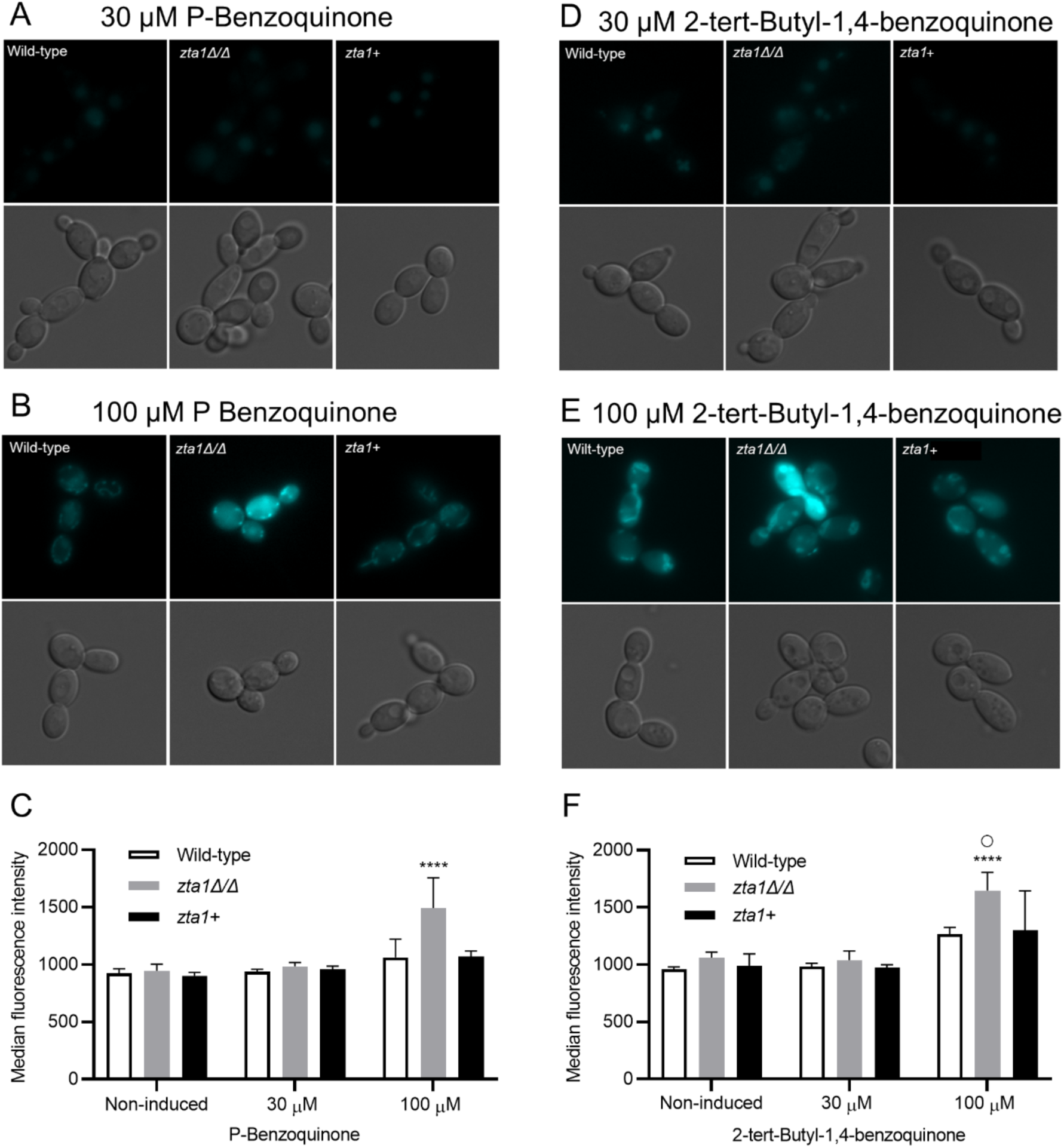
ROS accumulation is higher in the *zta1Δ/Δ* mutant exposed to quinones. Wild-type, *zta1Δ/Δ* and *zta1+* complemented strains were exposed to (A) 30 µM or (B) 100 µM p-benzoquinone, or (D) 30 µM or (E) 100 µM 2-tert-butyl-1,4-benzoquinone for 1h and then the accumulation of reactive oxygen species was assayed by incubating the cells with 2’,7’- Dichlorofluorescein diacetate for 20 min before imaging by fluorescence microscopy. (C) Median fluorescence intensity for panels A and B. (F) Median fluorescence intensity for panels D and E. Bars represent the average median fluorescence intensity of 3 independent assays on different days. ****, *P≤* 0.0001, by one-way analysis of variance (ANOVA). ○, *P≤* 0.05, by t-test (100 µM of p-benzoquinone vs. 100 µM 2-tert-butyl-1,4-benzoquinone) The wild type control strain was LLF100. The strains are described in Table 1.

**Table 1.** Strains used in this study.

| Strain | Parent | Genotype |
| --- | --- | --- |
| SN152 | SC5314 | *arg4Δ/ arg4Δ leu2Δ/leuΔ his1Δ/his1Δ URA3/ura3Δ::imm IRO1/iro1Δ::imm* |
| LLF100 | SN152 | *(Prototrophic wild-type control) ARG4/arg4Δ LEU2/leu2Δ HIS1/his1Δ URA3/ura3Δ::imm IRO1/iro1Δ::imm* |
| LLF054 | SN152 | *pst3-ycp4Δ::HIS1/pst3-ycp4Δ::LEU2 pst2Δ::frt/pst2Δ::frt pst1Δ::frt/pst1Δ::frt arg4Δ /arg4Δ* |
| LLF060 | LLF054 | *(Δ/Δ/Δ/Δ) pst3-ycp4Δ::HIS1/pst3-ycp4Δ::LEU2 pst2Δ::frt/pst2Δ::frt pst1Δ::frt/pst1Δ::frt ARG4/arg4Δ* |
| *zta1Δ/Δ* | SN152 | *zta1Δ::HIS1/zta1Δ::HIS1 ARG4/arg4Δ LEU2/leu2Δ URA3/ura3Δ::imm IRO1/iro1Δ::imm* |
| *zta1+* | SN152 | *(+ZTA1) zta1Δ::HIS1/zta1Δ::HIS1 ZTA1::NAT1::NEUT5L ARG4/arg4Δ LEU2/leu2Δ URA3/ura3Δ::imm IRO1/iro1Δ::imm* |
| Q Mut | LLF054 | *pst3-ycp4Δ::HIS1/pst3-ycp4Δ::LEU2 pst2Δ::frt/pst2Δ::frt pst1Δ::frt/pst1Δ::frt ARG4/arg4Δ* |
| Q Mut+ | LLF054 | *(+ZTA1) zta1Δ::HIS1/zta1Δ::HIS1 ZTA1::NAT1::NEUT5L pst3-ycp4Δ::HIS1/pst3-ycp4Δ::LEU2 pst2Δ::frt/pst2Δ::frt pst1Δ::frt/pst1Δ::frt ARG4/arg4Δ* |

### Zta1 acts in combination with the FLPs to protect *C. albicans* from quinones

To determine whether Zta1 is important for *C. albicans* resistance to quinones, we compared a wild-type *C. albicans* control strain with the *zta1Δ/Δ* mutant and the complemented strain *zta1+*. However, no differences in sensitivity were detected with disk diffusion halo assays when the strains were exposed to BZQ, MEN or TBBQ. Since *C. albicans* also possess QORs that belong to the FLP family, we tested whether *ZTA1* might be more important in the absence of the FLPs. *ZTA1* was deleted from a quadruple mutant lacking all four FLPs (*pst1Δ/Δ pst2Δ/Δ pst3Δ/Δ ycp4Δ/Δ*; referred to as Δ/Δ/Δ/Δ for simplicity) (7) to create a quintuple mutant lacking the four FLPs and *ZTA1* (referred to as Q Mut). We also generated a Q mutant complemented with *ZTA1* (*pst1Δ/Δ pst2Δ/Δ pst3Δ/Δ ycp4Δ/Δ zta1Δ/Δ + ZTA1*; referred to as Q Mut+). Although no difference was observed when cells were treated with BZQ or MEN, the Q Mut was more sensitive to killing by TBBQ when compared with the Δ/Δ/Δ/Δ mutant, and this sensitivity was partially reverted in the Q Mut+ complemented with *ZTA1* (Fig. 5). Perhaps TBBQ is more effective at killing *C. albicans* because it is more nonpolar than the other quinones used in this study and may therefore cross membranes more efficiently. These data indicate that Zta1 synergizes with the FLPs to detoxify certain quinones.

**Fig. 5.**
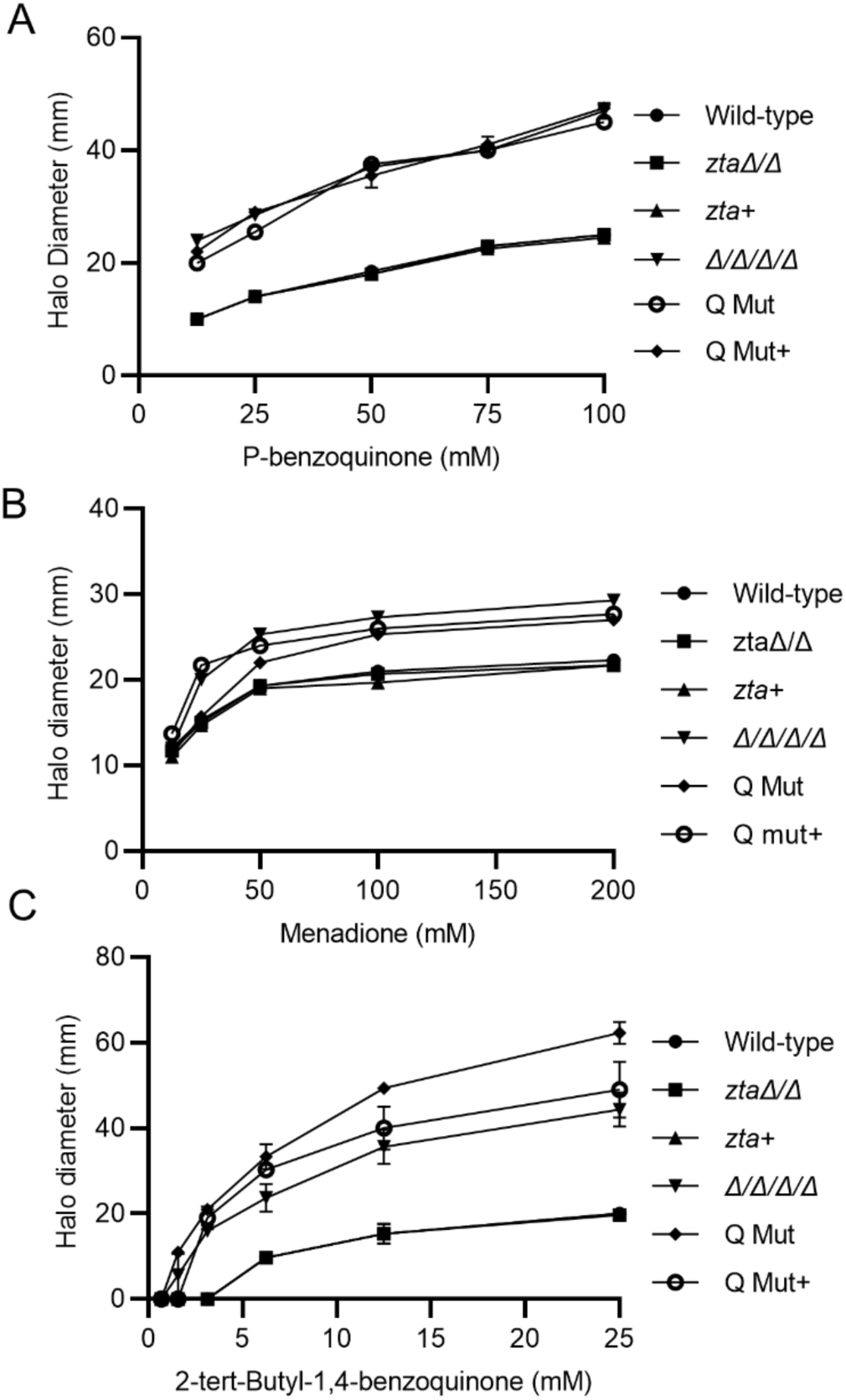
Zta1 acts in combination with FLPs to promote resistance to 2-tert-butyl-1,4- benzoquinone. Quantification of disk diffusion halo assays comparing the susceptibility of *C. albicans* strains to (A) p-benzoquinone, (B) menadione and (C) 2-tert-butyl-1,4-benzoquinone. The x-axis indicates the concentration of the compound applied to the disk and the y-axis indicates the diameter of the zone of growth inhibition. The strains tested included the wild-type control strain (LLF100), *zta1Δ/Δ*, *zta1+*, *Δ/Δ/Δ/Δ*, Q Mut, and Q Mut+ complemented strain (see Table 1). Results represent the averages from at least three independent experiments, each done in duplicate. Error bars indicate SD.

### Zta1 protection appears to be specific to quinones

To determine if Zta1 protects *C. albicans* against other sources of oxidative stress, the susceptibility of cells to other oxidizing agents was examined including H_2_O_2_, tert-Butyl-hydroperoxide (Fig. 6) and the thiol oxidizing compound diamide (data not shown). Neither *zta1Δ/Δ* nor the Q Mut strain showed increased susceptibility to these other oxidizing agents (Figure 6). Thus, although Zta1 is induced by H_2_O_2_, it does not appear to play a major role in protecting against this type of oxidative stress.

**Fig.6.**
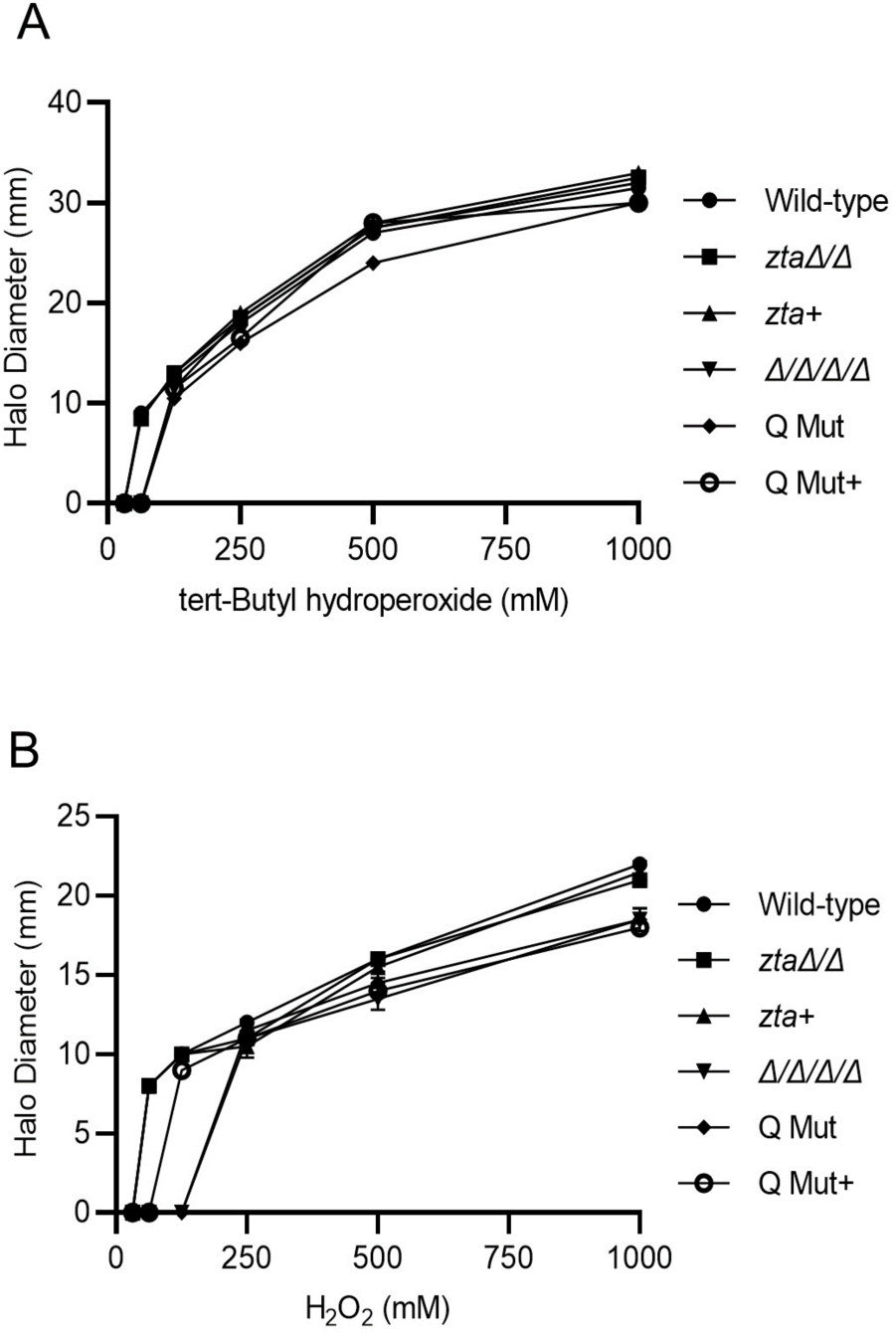
The *zta1*Δ/Δ mutant does not show increased susceptibility to peroxides. Quantification of disk diffusion halo assays comparing the susceptibility of *C. albicans* strains to tert-butyl-hydroperoxide and hydrogen peroxide (H_2_O_2_). The strains tested included the wild-type control strain (LLF100), *zta1Δ/Δ*, *zta1+, Δ/Δ/Δ/Δ*, Q Mut, and Q Mut+ complemented strain (see Table 1). Results represent the averages from at least three independent experiments, each done in duplicate. Error bars indicate SD.

### *zta1*Δ/Δ shows a trend of increased susceptibility to killing by neutrophils

To assess the potential significance of *ZTA1* in *C. albicans* resistance to attack by the immune system, we analyzed the ability of human neutrophils to kill *C. albicans* wild-type and mutant strains. Notably, increased vulnerability of *zta1Δ/Δ* cells to neutrophil-mediated killing was observed, which was reversed in the *ZTA1* complemented strain (Fig. 7). Unfortunately, due to variation in the data, this trend did not reach statistical significance when analyzed by ANOVA or t-test. Donor-dependent variations when working with neutrophils are known to occur, since environmental aspects such as temperature, pH, oxygen and glucose levels can have a strong influence in the function of neutrophils (22). Deletion of the *ZTA1* gene in the Δ/Δ/Δ/Δ mutant did not yield an obvious increase in susceptibility to neutrophils; similar survival levels were observed between the *zta1Δ/Δ*, Δ/Δ/Δ/Δ, Q mut and Q mut+ (complemented with *ZTA1*).

**Fig. 7.**
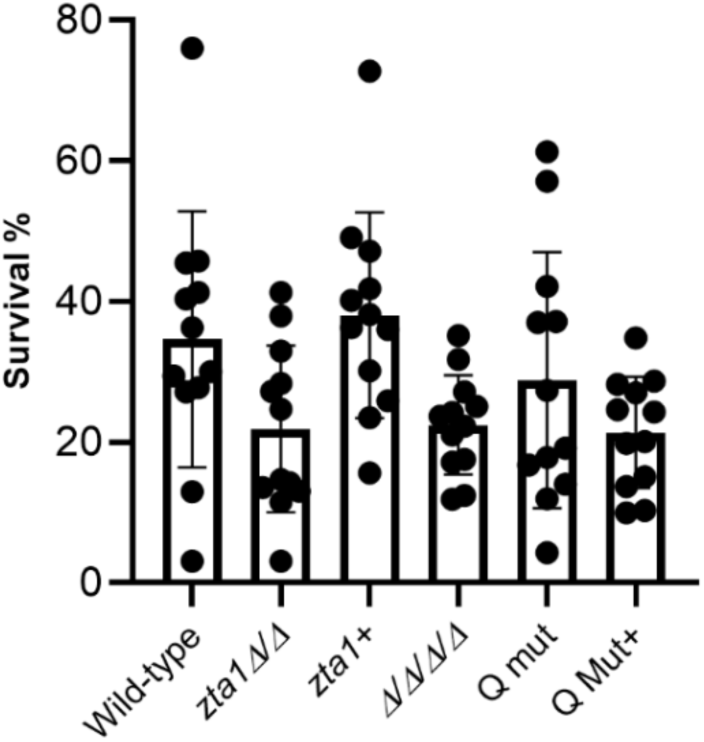
Susceptibility of C. albicans strains to attack by human neutrophils. Human neutrophils from 4 different donors were incubated with the wild-type control strain (LLF100), *zta1*Δ/Δ, *zta1+*, FLP mutant (Δ/Δ/Δ/Δ/), Q Mut and Q Mut+ (complemented with *zta1*) during 4 h. Yeast viability and percentage of survival was determined using PrestoBlue dye. Data are presented as the mean of four independent experiments, each done in triplicate. Error bars indicate SD. Strains are described in Table 1.

### *ZTA1* role in *C. albicans* virulence

The role of *ZTA1* in *C. albicans* virulence was investigated in a mouse model of hematogenously disseminated candidiasis (23). The wild-type, Δ/Δ/Δ/Δ, Q Mut and Q Mut+ strains were used infect BALB/c mice via tail vein injection with 2.5 x 10^5^ *C. albicans* cells. Since it has been shown that the Δ/Δ/Δ/Δ mutant is non-virulent and is cleared after ∼4 days of infection (7), we focused on measuring the colony forming units in the kidney (CFU/g kidney) after 2 and 3 days of infection. The kidneys are an established target organ for testing the capacity of *C. albicans* to cause an infection, since growth occurs rapidly in the kidneys during the first 2 days after injection (24). Interestingly, at day 2 (Fig. 8A) and 3 (Fig. 8B) post infection, the Q Mut strain showed lower CFU/g kidney than did the Δ/Δ/Δ/Δ strain, suggesting a role for *ZTA1* in virulence. In contrast, the Q Mut+ strain had similar fungal burden as the Δ/Δ/Δ/Δ, indicating this effect was reversed by reintroducing the *ZTA1* gene. However, this trend in the results did not reach statistical significance when analyzed by ANOVA or t-test. Nonetheless, the *Δ/Δ/Δ/Δ* and Q Mut strains had lower kidney fungal burden than the wild-type control strain, highlighting the importance of quinone reductases in virulence. Although statistical significance was lacking, the overall trends of these studies suggest that Zta1 contributes to *C. albicans* virulence.

**Fig. 8.**
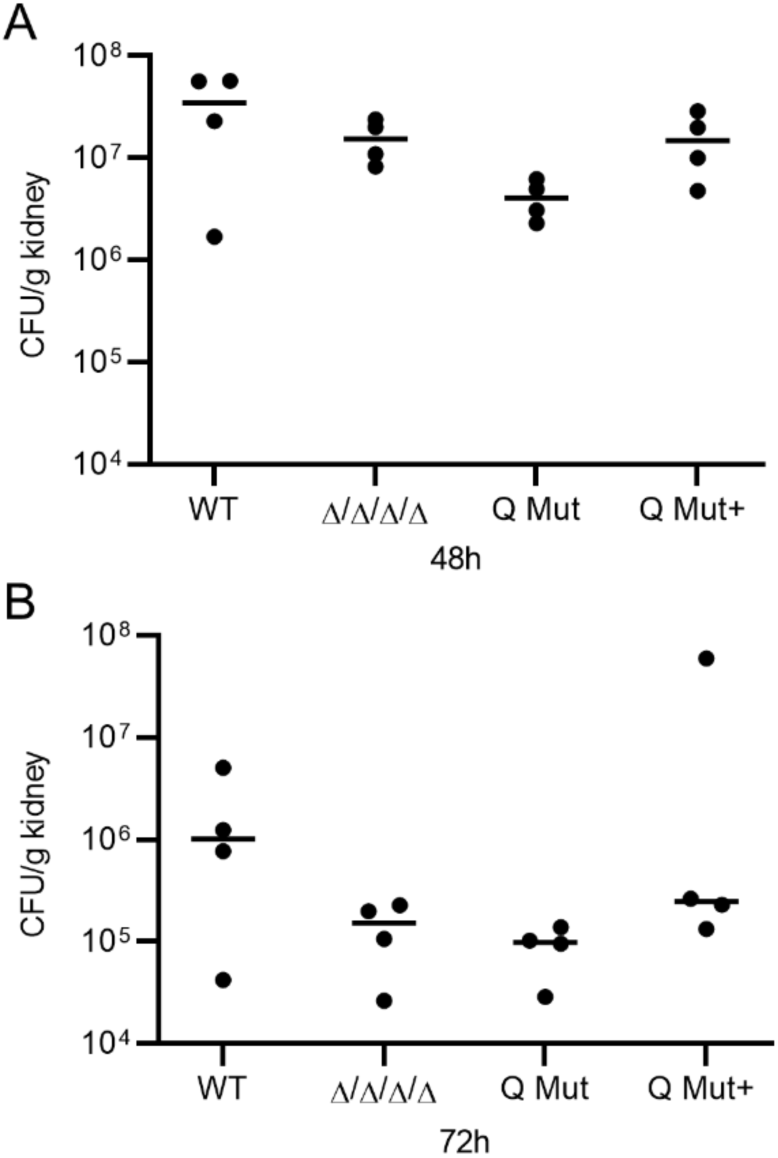
Zta1 role in *C. albicans* virulence. Female Balb/c mice were infected with 2.5 x 10^5^ cells of the indicated *C. albicans* strain via the tail vein. Strains used included the LLF100 wild-type control strain, FLP mutant (Δ/Δ/Δ/Δ/), Q Mut and Q Mut+ (complemented with *zta1*) (see Table 1). CFU/g of kidney was determined at (A) day 2 and (B) day 3 post infection. Four mice were injected with each strain for each day of analysis and their mean is presented.

## DISCUSSION

Fungal cells face various challenges when colonizing a host, including reactive oxygen species (ROS) that can cause widespread damage to proteins, lipids, and nucleic acids (25). Among the arsenal employed by fungal cells to deal with this stress are quinone oxidoreductase (QOR) enzymes capable of reducing quinones into hydroquinones (13). This mechanism holds significance as quinones have the potential to alkylate proteins and DNA or undergo autoxidation, resulting in the production of semiquinone radicals and the generation of ROS (9, 11, 26, 27). Furthermore, QORs can also reduce ubiquinone, a well-known component of the mitochondrial membrane that participates in cellular respiration, but is also present in other membranes, including the plasma membrane, where it functions as an antioxidant (28). In fact, recent studies revealed that four members of the FLP family of flavin-dependent NADPH:quinone oxidoreductases (Pst1, Pst2, Pst3, Ycp4) are needed for resistance to quinones and virulence (7). The FLP family members in C. albicans are thought to act in part by catalyzing a two-electron reduction of ubiquinone that enable it to then reduce ROS in the plasma membrane (7). The FLPs are localized to specialized plasma membrane domains called eisosomes, which is thought to facilitate a role in protecting the *C. albicans* plasma membrane polyunsaturated fatty acids (PUFAs) which are more prone to lipid peroxidation. This type of lipid oxidation can trigger a chain reaction that spreads to other PUFAs and then the peroxidized lipids can damage proteins and DNA (8, 29). Another QOR present in fungi is the ζ-crystallin-like protein Zta1, which belongs to a structurally distinct family that is believed to catalyze a one-electron reduction of quinones (16, 18). Due to the FLPs importance, we decided to investigate the role of Zta1 in *C. albicans*.

The high degree of amino acid conservation in the active site of *C. albicans* Zta1 strongly indicates that it functions as a QOR, as in *S. cerevisiae* (Fig. 1). Supporting this notion, *C. albicans* Zta1 was rapidly induced upon exposure to p-benzoquinone (BZQ), 2-tert-butyl-1,4- benzoquinone (TBBQ), menadione (MEN) and hydrogen peroxide (H_2_O_2_) (Figs. 2 and 3). However, despite its induction by H_2_O_2_, Zta1 does not appear to play a significant role in protecting cells against H_2_O_2_ or other oxidants such as tert-butyl-hydroperoxide (Fig. 6) and diamide (data not shown). Previous studies have reported the upregulation of flavin-containing QORs under certain conditions, such as increased temperature (26). Moreover, Zta1 production in *S. cerevisiae* increased following treatment with rapamycin, heat shock and during the stationary phase, when cells are starved, and toxic compounds accumulate (30). These observations suggest that Zta1 may be part of a more general fungal cellular response to stress, which could explain its induction by H_2_O_2_ while not directly providing protection against it.

Assays utilizing H_2_DCFDA revealed that the *zta1Δ/Δ* exhibited increased accumulation of ROS upon exposure to quinones, indicating it protects against this type of oxidative stress (Fig. 4). Surprisingly, we did not observe increased cell death of the *zta1Δ/Δ* mutant in response to quinones, despite a previous report that the *S. cerevisiae zta1Δ* mutant showed slightly increased susceptibility to menadione (18). However, the Q Mut, which lacks all four FLPs (*pst1Δ/pst2Δ/pst3Δ/ycp4Δ*) as well as *ZTA1,* exhibited greater susceptibility to TBBQ, although not to BZQ or MEN. One possibility for this specific susceptibility is that TBBQ, being the most non-polar quinone tested in our study (8), may cross the plasma membrane more readily and enter the cytoplasm where Zta1 is located (Fig. 2). In this scenario, *C. albicans* FLPs may preferentially act at the plasma membrane where they are localized and Zta1 may be important to reduce TBBQ in the cytoplasm, where it was detected.

To determine if *ZTA1* is important for *C. albicans* to avoid attack by the immune system, the *zta1ΔΔ* mutant was assessed for its ability to survive incubation with human neutrophils. The results showed a trend of increased killing of the *zta1ΔΔ* mutant by neutrophils, which was reversed by reintroduction of the *ZTA1* gene in the complemented strain (Fig. 7). The presence or absence of *ZTA1* in cells lacking all four FLPs did not appear to obviously impact in *C. albicans* survival, perhaps because cells lacking the FLPs are already more susceptible to neutrophils.

Previous studies demonstrated that the FLPs are critical for *C. albicans* infection in mice, as a mutant lacking all four FLPs was avirulent and cleared from the kidney at early times after infection that correlated with the influx of neutrophils (7). Considering the limited role of *zta1Δ/Δ* in susceptibility to quinones in vitro, we decided to investigate whether *zta1Δ/Δ* would exacerbate the previously described virulence defect of the FLP mutant (*pst1Δ/Δ pst2Δ/Δ pst3Δ/Δ ycp4Δ/Δ*) (7) by assessing the kidney fungal burden in infected mice. Notably, we observed a trend towards reduced CFU/g kidney of the Q Mut at both day 2 and day 3 post-infection. Complementation of Q Mut with *ZTA1* restored its clearance to levels comparable to those of the FLP mutant. This finding aligns with studies implicating QORs in the virulence of other organisms, such as *Xanthomonas citri*, *Mycobacterium turbeculosis* and *Staphylococcus aureus* (31-33). This suggests that Zta1 contributes, at least partially, to *C. albicans* ability to resist in the host and evade its immune system.

Collectively, our data indicate that Zta1 is rapidly induced by quinones and protects *C. albicans* against quinone-induced damage. It also protects against the accumulation of ROS caused by quinones and might contribute to *C. albicans* capacity to establish an infection. Zta1 function seems to overlap with other QORs, such as the FLPs, and due to their difference in localization, they seem to complement each other. These results highlight the important role of QORs in *C. albicans* repertoire of strategies to be a successful pathogen.

## METHODS

### Strains and media

*C. albicans* strains used are listed in Table 1. Strains were kept in YPD (1% yeast extract, 2% peptone, 2% glucose) agar plates. Prior to experiments, cultures were grown in YPD medium (2% dextrose, 1% peptone, 2% yeast extract, 80 mg/L uridine) (34). A 3xGFP*γ* tag was fused to the 3’ end of the open reading frame of *ZTA1* by homologous recombination as previously described (35, 36). The DNA was introduced into *C. albicans* cells by electroporation and allowed to recombine with the homologous regions of the *ZTA1* gene. Strains were verified by PCR analysis and microscopic examination of GFP*γ* fluorescence in a Zeiss Axiovert 200M microscope equipped with an AxioCam HRm camera and Zeiss ZEN software. Mutant strains of *zta1* were generated using a transient expression of CRISPR-Cas9 to obtain homozygous deletion of the target gene, as previously described (37). A 20-bp target sequence of the sgRNA was used to delete *ZTA1* in strain SN152, and primers were designed to include 80 bases of homology to the sequences of target gene. The gene was deleted with by replacement with a *HIS1* selectable marker. For the Q Mut, *ZTA1* was deleted and replaced with an *ARG4* selectable marker in strain LLF054. The complemented strains of *ZTA1* were obtained by integrating a copy of the wild-type gene sequence into the *NEUT5L* region of the genome using the gap-repair method, as previously described (38). The PCR fragment was constructed by amplification of the genomic DNA from 0.5kb upstream of the start codon and 0.25kb downstream of the stop codon of the *ZTA1* gene from *C. albicans* SC5314 using primers 5227 and 5228, which includes 20-bp homology to the *ZTA1* upstream and downstream regions and 40-bp homology with the plasmid pDIS3, that carries the *NAT1* resistance cassette. The PCR fragment was recombined into *Sma*I-digested pDIS3 in *S. cerevisiae* strain L40, generating the pDIS3-*ZTA1-NAT1* construct, which was subsequently released from the pDIS3 vector by *Sfi*I digestion and transformed into the *zta1Δ/Δ* or Q Mut strain by electroporation. The primers used are listed on Table 2.

**Table 2.**
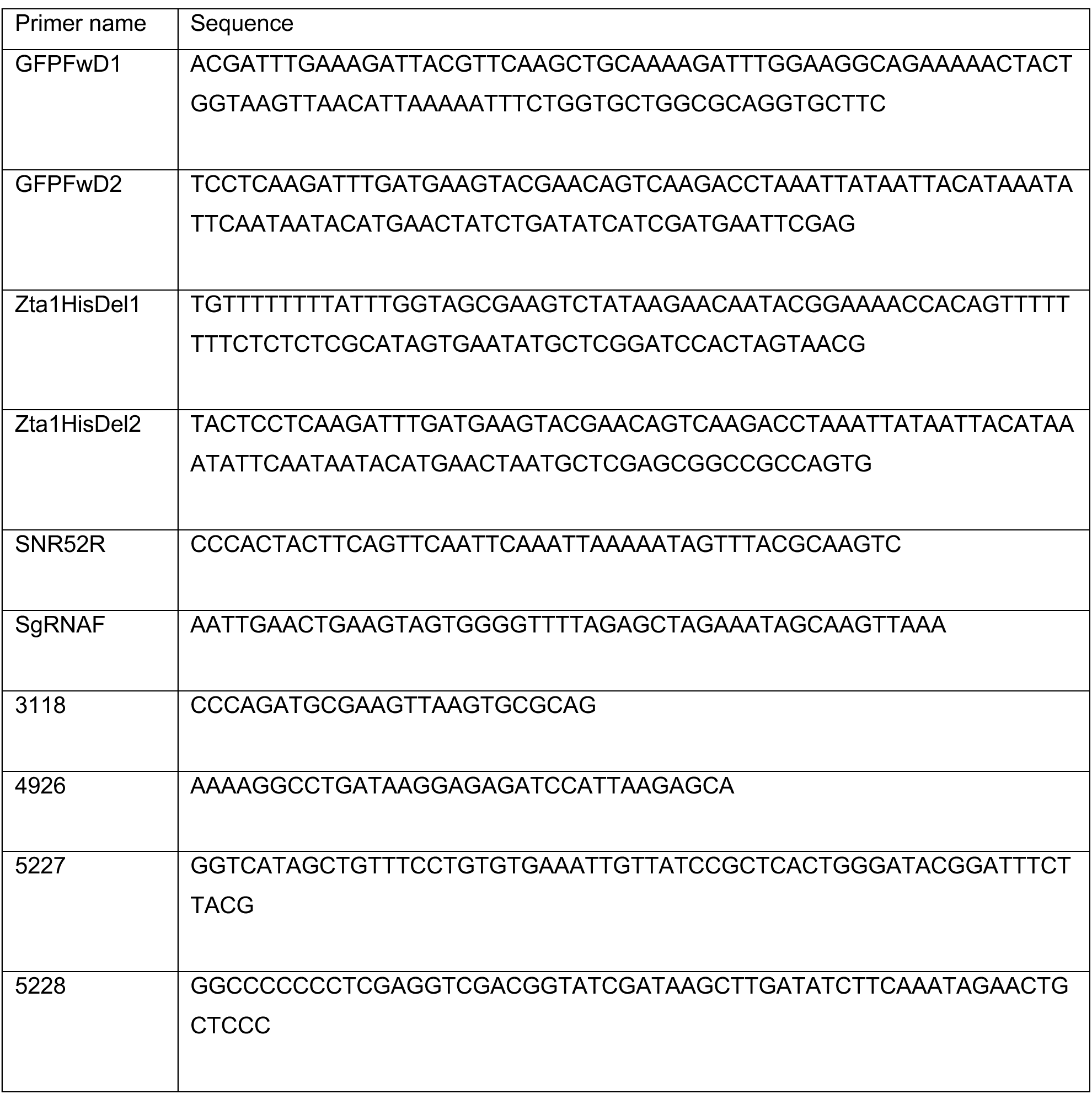
Primers used in this study.

### Sequence alignments of Zta1 proteins

Zta1 protein sequence was obtained from the *S. cerevisiae* Genome Database (http://www.yeastgenome.org). BLAST searches were carried out to identify Zta1 homologues in the *C. albicans* Genome Database (http://www.candidagenome.org) and genome sequences present at the Web site of the National Center for Biotechnology (http://www.ncbi.nlm.nih.gov/BLAST/). Multiple sequence alignments of the predicted Zta1 proteins were carried out using Jalview (39).

### Microscopic analysis of Zta1-3xGPF*γ* production

3xGFP*γ* tagged *C. albicans* strains were grown overnight in YPD at 30°C and then diluted 1:250 in 5 mL YPD and grown until a density of about 1 × 10^7^ cells/ml. Cells (1 mL) were treated with 100 µM of one of the quinones p-benzoquinone (BZQ; Sigma-Aldrich, St. Louis, MO), 2-tert-butyl-1,4-benzoquinone (TBBQ; Cayman Chemical, Ann Arbor, MI), menadione (MND; Sigma-Aldrich, St. Louis, MO) or 500 µM of H_2_O_2_ and incubated for 1h at 30°C on a tube roller. Samples were centrifuged, washed in sterile phosphate-buffered saline (PBS), and analyzed by fluorescence microscopy. To assess the effect on Zta1 production of different times and concentrations of incubation with BZQ, cells were treated with 10, 30 or 100 µM of BZQ, or treated with 100 µM of BZQ for 15, 30 and 60 minutes of incubation and then prepared for imaging as described above. Zeiss ZEN software was used to control the microscope and for deconvoluting images and calculating the mean fluorescence intensity (MFI) of cells. Statistical analysis of MFI compared to non-treated GFP-tagged cells was carried out with Prism 6 software (GraphPad Software, Inc., La Jolla, CA).

### Western blot analysis of Zta1-3xGFP*γ* levels

The same protocol described above was used for western blot analysis. Shortly, 1 × 10^7^ cells/ml were treated with 100 µM of BZQ, TBBQ, MEN or 500 µM of H_2_O_2_ and incubated for 1h at 30°C shaking before being harvested. The same process was used for cells treated with 10, 30 or 100 µM of BZQ for 1h or cells treated with 100 µM of BZQ for 15, 30 or 60 minutes. Cell lysates were prepared by bead-bashing as follows: 20 mL of cell culture prepared as described above were centrifuged, washed with PBS and lysed using 300 µl of 1x Laemmli buffer (2% SDS, 10% glycerol, 125 mM Tris-HCl, pH 6.8, and 0.002% bromophenol blue, 5% 2- Mercaptoethanol) and zirconia beads by 5 rounds of 1 min of bead beating followed by 1 min on ice. Samples (10µl) were separated by SDS-PAGE and transferred to a 0.4 mm nitrocellulose membrane using a semidry transfer apparatus. Blots were probed for 1h at room temperature (RT) with a mouse anti-GFP antibody (Living Colors -JL-8, BD Biosciences Clontech) in TBS-T buffer (20 mM Tris, 150 mM NaCl, 0.1% Tween 20, 2% [wt/vol] bovine serum albumin [BSA], and 0.2% [wt/vol] sodium azide). The blots were then washed and incubated for 1h with an IRDye 800-conjugated anti-mouse IgG (LI-COR Biosciences, Lincoln, NE) diluted 1:15,000 in TBS (20 mM Tris, 150 mM NaCl) containing 0.5% Tween 20. Blots were washed using TBS-T and visualized by scanning with an Odyssey CLx infrared imaging system (LI-COR Biosciences). The resulting images were analyzed using Image Studio software (LI-COR Biosciences). For Coomassie-stained gels, SDS-PAGE was performed as described above, and then gels were stained in a Coomassie brilliant blue solution (0.1% Coomassie R-250, 40% ethanol, 10% glacial acetic acid) overnight. Gels were destained in destaining solution (40% methanol, 10% acetic acid) and analyzed using Image Studio software (LI-COR Biosciences). Statistical analysis of densitometry was carried out with Prism 6 software (GraphPad Software, Inc., La Jolla, CA) using multiple *t* tests against non-treated GFP-tagged cells.

### Sensitivity to oxidizing agents

*C. albicans* strains were grown overnight in YPD at 30°C were diluted to 2.5 x 10^5^ cells were spread onto the surface of a synthetic complete medium agar plate. 10 µl of each compound was applied to paper filter disks (Becton, Dickinson and Company, Sparks, MD), the disks were applied to the plates surface, incubated at 30°C for 48 h, and then the diameter of the zone of growth inhibition (halo) was measured. Compounds tested included p-benzoquinone (BZQ; Sigma-Aldrich, St. Louis, MO), menadione (MND; Sigma-Aldrich, St. Louis, MO), 2-tert-butyl-1,4-benzoquinone (TBBQ; Cayman Chemical, Ann Arbor, MI), hydrogen peroxide (H_2_O_2_; Sigma-Aldrich, St. Louis, MO), tert-Butyl hydroperoxide (tBOOH – Acros Organics) and diamide (Sigma-Aldrich, St. Louis, MO). For the CFU assays, wild-type and the 3xGFP-tagged strains were grown overnight, harvested by centrifugations, and resuspended in PBS. A total of 1 × 10^7^ cells/ml were inoculated in liquid YPD and treated with one of the following: 100 µM of p-benzoquinone, 100 µM 2-tert-butyl-1,4-benzoquinone, 100 µM menadione or 500 µM of hydrogen peroxide. Tubes were incubated for 1 h at 30°C on a tube roller, after which cultures were centrifugated and cells washed twice with PBS. Serial dilutions were plated in YPD plates and incubated for 48h at 30°C, and colony forming units were counted.

### ROS accumulation

*C. albicans* (WT, *zta1Δ/Δ* and *zta1+*) was grown overnight in 5 mL of YPD. The following day. cultures were diluted to 0.100 OD into 5 ml of fresh YPD and grown for 2h at 30°C agitating. BZQ or TBBQ (10, 30 and 100 µM/mL) were added, and cultures incubated for an additional 1 h. After that, H_2_DCFDA (10 µM/mL) was added, and cultures were incubated for 20 minutes in the dark at 30°C with shaking. Cultures were centrifuged, washed twice with PBS and resuspended in 100µL of PBS. Samples were analyzed by microscopy at 492nm in a Zeiss Axiovert 200M microscope equipped with an AxioCam HRm camera and Zeiss ZEN software for deconvolving images.

### Human neutrophil killing of *C. albicans*

Blood was obtained from study participants with written informed consent through a protocol approved by the University of Wisconsin Internal Review Board. Neutrophils were isolated from 4 different donors using the MACSxpress Neutrophil Isolation and MACSxpress Erythrocyte Depletion kits (Miltenyi Biotec Inc., Auburn, CA) and suspended in RPMI 1640 (without phenol red) supplemented with 2% heat-inactivated fetal bovine serum (FBS) and supplemented with glutamine (0.3 mg/ml) as previously described (40). For the killing assays, neutrophils (4×10^5^ cells) and the *C. albicans* strains (1×10^6^ cells) were added to wells of a culture treated 96-well flat-bottom plate and incubated for 4 h at 37°C in 5% CO_2_. After incubation, 10 µg of DNAse1 was added to each well and the plate was incubated at 37°C with 5% CO_2_ for 10 minutes, after which contents of the wells (media and non-adherent cells) were moved into a 96-well round-bottom plate. Some neutrophils and *C. albicans* cells can adhere to the flat-bottom plate, so the contents of both the flat-bottom plate and the round-bottom plate were processed and combined back in the flat-bottom plate for the analysis. The round-bottom plate was centrifuged at 1,200xg for 2 min, the supernatant was discarded and 150 µl of ddH_2_O containing 1 µg/ml DNAse1 was added to each well of the round-bottom and the flat-bottom plate. Plates were incubated at 37°C for 20 minutes to lyse neutrophils. The round-bottom plate was centrifuged at 1,200xg and the supernatant removed, after which the contents of the original flat-bottom plate were removed and used to resuspend the contents of their respective wells in the round-bottom plate. The lysis step was repeated one additional time, after which the round-bottom plate was centrifuged at 1,200xg, the supernatant removed, and the remaining yeast cells were resuspended in 100 µl of RPMI + 2% FBS and transferred to the original wells of the 96-well flat-bottom plate. Then, 10 µl PrestoBlue (Invitrogen, Eugene, OR) was added to each well, gently mixed, and plate was incubated for 25 min at 37°C. Fluorescence 560/590 was then read on a BioTek Synergy|H1 microplate reader (Agilent, Santa Clara, CA). Percentage of viable *C. albicans* cells was quantified by calculating the fluorescence signal of the treated well (mutant and neutrophils) as a percentage of the control well (same mutant and no neutrophils).

### Mouse infection assays

Fungal burden was tested in mice as previously described (24) using a protocol approved by the Stony Brook University IACUC committee. Strains were grown overnight in YPD medium, reinoculated into fresh medium and incubated again overnight. Cells were harvested after centrifugation and washed twice with PBS and counted using a hemocytometer. Cells were diluted to 1.25 x 10^6^ cells/ml with PBS. Female BALB/c mice (8 weeks old) were injected via the lateral tail vein with 2.5 x10^5^ cells (200µl). After 48h or 72h, kidneys were excised, weighed, and then homogenized in 5 ml PBS for 30 s with a tissue homogenizer (Pro Scientific Inc.). The CFU per gram of kidney (CFU/g kidney) was determined by plating dilutions of the homogenates on YPD agar medium plates and incubating for 2 days at 30°C. Statistical analysis of the CFU data was carried out with Prism 6 software (GraphPad Software, Inc., La Jolla, CA) using one-way analysis of variance with one-way ANOVA.

## ACKNOWLEDGMENTS

We thank the members of our lab for their helpful advice and suggestions on this study.

This research was supported by Public Health Service grants awarded from the National Institute of Health to J.B.K (R01AI047837) and to J.E.N (R01AI145939 and R21AI159583).

